# Contribution of the Type 3 Secretion System to immunogenicity of a live *Yersinia pseudotuberculosis* plague vaccine

**DOI:** 10.1101/2024.06.11.598433

**Authors:** Anne Derbise, Chloé Guillas, Hebert Echenique Rivera, Elisabeth Carniel, Chris Gerke, Javier Pizarro Cerdá, Christian E. Demeure

## Abstract

The causative agent of plague, *Yersinia pestis*, remains a treat for public health worldwide. In the perspective to develop effective and safe vaccines, we present here a derived version of our *Y. pseudotuberculosis* VTnF1 live attenuated vaccine candidate that lacks the pYV virulence plasmid coding for the Type 3 Secretion system (T3SS) and no antibiotic resistance cassettes. This strain, called VpYV-, was safe for immunodeficient mice, and thus can be considered as deeply attenuated. It still has tropism for lymphatic tissues (Peyer’s patches, Mesenteric lymph nodes) but hardly reaches the spleen and liver. It elicits IgG to the F1 antigen as efficiently as VTnF1 but less directed to other Yersinia antigens. A single oral dose induced 100% protection against bubonic and pneumonic forms of plague, but this protection decreased faster with time than that of VTnF1. In the same line, VpYV-was 30% less protective against a F1-negative *Y. pestis*, revealing that the tools encoded by pYV are mandatory to obtain a large spectrum protection. Finally, VTnF1 like the *Y. pestis* vaccine EV76 can induce protection against co-infected *Y. pestis* relying on ‘host iron nutritional immunity”, indicating a potential use as therapeutics of recent infection. In contrast VpYV-failed to do so, revealing an importance of the T3SS in this mechanism. Overall, VpYV- and its parental strain VTnF1 offer a choice between more attenuation and safety or vaccine performances. They give an alternative and may represent useful tools to prevent and treat *Y. pestis* infection in healthy or immunosuppressed individuals.

## Introduction

*Yersinia pestis*, the terrible causative agent of plague, is among the deadliest bacteria affecting humans. It is a Gram-negative bacillus derived very recently from the much less virulent enteropathogen *Yersinia pseudotuberculosis* [1]. Following a limited number of genetic changes, *Y. pestis* adopted a new lifecycle using fleas as vector and rodents as reservoir. *Y. pestis* is therefore mainly a zoonotic pathogen and is present in animals in large endemic territories throughout the world. For this reason, plague cannot be eradicated and will always have a capacity to pass from its animal reservoir to humans to cause outbreaks. Despite considerable progress in prevention and cure, natural foci still exist in Africa, Asia and the Americas, and plague is endemic in more than 25 countries worldwide, including China, the USA, Peru, the Democratic Republic of Congo, and India. Madagascar is the most affected country worldwide, with hundreds of cases every year. Since the beginning of the nineties, plague is categorized as a re-emerging disease [2]. Also, due to the bioterrorism threat, *Y. pestis* is classified by the USA Center for Disease Control (CDC) among the Tier 1 select biological agents that pose a risk to national security [3].

The plague bacillus is generally transmitted to humans via the bite of an infected flea in skin. It causes bubonic plague, the most frequent clinical form of the disease which is lethal in 50-70% of patients if untreated. Its clinical signature is the development of a painful infected lymph node (bubo), with high fever and a severe faintness. *Y. pestis* then spreads systematically via blood and infection is lethal in 50-70% of patients if untreated. If infection reaches the airways, it becomes highly contagious due to the emission of infected aerosols responsible for inter-human transmission of the bacilli. The resulting pneumopathy is systematically lethal in usually less than three days.

*Y. pestis* is sensitive to most antibiotics which are the main therapy against the ongoing infection. Despite this, the fast development of the disease requires a fast access to medical help, and plague lethality in endemic regions remains at 10% or more. Moreover, the appearance in Madagascar of *Y. pestis* strains showing high-level resistance to several antibiotics currently used to treat patients, has evidenced the limits of current therapies [4].

Because plague would be difficult to eradicate from its animal reservoirs and antibiotics could fail to offer full therapeutic coverage, vaccines might represent an alternative to limit the death toll in humans. With its prophylactic mode of action, vaccination might limit the emergence of plague outbreaks, however, no plague vaccine is presently marketed in occidental countries.

We previously proposed a vaccine strategy against plague consisting in oral vaccination with a live, attenuated *Y. pseudotuberculosis* [5], the genetically very close parent of *Y. pestis*. Whereas the *Y. pestis* genome is instable, *Y. pseudotuberculosis* has a much higher genomic stability, and is also much less pathogenic. The *Y. pseudotuberculosis* IP32953 strain was irreversibly attenuated by deletion of genes encoding three essential virulence factors. These are the High pathogenicity island (HPI, a very efficient iron capture system), the adhesin PsaA, and the YopK toxin. To additionally increase vaccine efficiency, the *Y. pestis caf* operon was inserted in the chromosome to induce the production of the very immunogenic F1 antigen forming the *Y. pestis* specific pseudocapsule, a protective target for immunity [6].

In all three Yersiniae species pathogenic for superior animals (*Y. pestis, Y. pseudotuberculosis* and *Y. enterocolitica*), curing the pYV plasmid (pCD1 in *Y. pestis*) results in a failure to sustain a high rate of multiplication *in vivo* and to cause disease [7], [8], [9]. This plasmid encodes the Type 3 Secretion System (T3SS), a key virulence factor which injects effector Yop toxins into host cells, resulting in a suppression of the innate immune response [8]. The adaptative immune response is also affected via effects on dendritic cells (DC), the main cells presenting antigen to naive T cells [10]. As a result, both the T4 and T8 responses are impaired. When building our VTnF1 strain, deletion of *yopK* was chosen because YopK blocks the flux of effector Yops [11], thus reducing the bacteria’s ability to cause systemic infection [12]. By doing so, all other T3SS antigens were still present to serve as immune targets, but their contribution to virulence was reduced. The VTnF1 strain induces antibodies against Yops among other antigens [13] but was highly attenuated by both oral (ig) and subcutaneous (sc) routes [14].

The efficiency of a live vaccine requires both immunogenicity and attenuation, and they often trigger powerful humoral and cell-mediated immune responses [15], [16]. However, immunocompromised subjects cannot be vaccinated with attenuated live vaccines because they fail to control bacteria multiplication and are at risk of a potentially harmful infection. For example, the historical live-attenuated *Y. pestis* vaccine EV76 [17] has a functional T3SS and is not completely avirulent. Payne *et al*. reported that EV76 induces mouse mortality when associated with cortisone or iron [18]. In contrast, pYV-cured strains did not become virulent in cortisone or iron-treated mice and were thus truly avirulent.

The attenuated *Y. pseudotuberculosis* VTnF1 strain triggers only moderate and transient clinical signs in immunocompetent laboratory mice but was previously not tested in immunocompromised animals. To potentially enhance our vaccine’s safety, we explored the possibility that curing of the pYV plasmid to fully eliminate the T3SS -including Yops-could attenuate the strain even more deeply without affecting its immunogenicity. It was previously reported that pYV-negative *Y. pseudotuberculosis* strains retain the ability to transiently disseminate to the spleen [19], [20], retain their tropism for lymphocyte-rich regions of lymph nodes and can induce a protective immune response [21].To this aim a VTnF1 sub-strain cured of the pYV plasmid was derived. It was renamed VpYV- and was compared to the original VTnF1 strain for its efficiency as plague vaccine.

## Materials and Methods

### Culture conditions and bacterial strain construction

*Y. pseudotuberculosis* and *Y. pestis* strains and derivatives (Supplementary Table S1) used were previously described [22]. *Y. pestis* was cultured on Luria-Bertani plates containing 0.002% hemin (LBH) for 48h at 28°C. Bacterial density was evaluated by spectrometry at 600 nm and numeration on LBH plates. Bacteria were grown at 28°C (for *Yersinia pseudotuberculosis* background strains) or 37°C (for *E. coli*) in Lysogeny broth (LB) or on LB agar (LBA) plates. Kanamycin (Km ; 30 μg/ml), trimethoprim (Tmp ; 100 μg/ml), chloramphenicol (Cm ; 25 μg/ml), spectinomycin (Spec ; 50 μg/ml), sucrose (10% weight/vol) or Irgasan (0,1 μg/ml) were added to the media when necessary. All experiments involving *Y. pestis* were performed in a BSL3 laboratory.

Cloning of the upstream and downstream *Yersinia*’s flanking regions of the Km, Tmp and Spec cassettes in the pCVD442 plasmid were performed using a standard cloning methodology where fragments were first PCR amplified using oligomers containing 5’ restriction sites extensions (Supplementary Table S2) and then subjected to enzymatic restrictions and ligation to pCVD442 according to manufacturer’s instructions. After selection of the correct recombinant pCVD442b, each plasmid was introduced by electroporation into the SM10 *λ*pir *E. coli* strain. SM10 recombinants were then used as donor strains to transfer pCVD442 recombinant plasmids by mating with the *Y. pseudotuberculosis* vaccine candidate strain following a protocol previously described[23]. This procedure allowed to generate a new version of the vaccine candidate named VTnF1-S, sensitive to all antibiotics. For comparison, the original strain containing the cassettes is hereafter named “VTnF1-R”.

DNA was extracted from the VTnF1-S vaccine strain, and the complete genome was sequenced (paired-end 150 bases, NextSeq500, Illumina). An assessment of modifications was obtained both by the *de novo* assembly of the genome and mapping of the reads against the original genome (strain IP32953, GenBank accession number: BX936398) to establish both chromosome and pYV plasmid maps using the CLC Assembly Cell 4.4.0 software (CLC bio, Waltham, USA). The genome maps were annotated, and new coding sequences identified.

The VTnF1-S strain was cured of the pYV plasmid by culture at 37°C in liquid LB medium with shaking prior to culture at 37°C on MOX-Congo plates (low calcium conditions) [24]. Plasmid-cured bacteria were then identified by their capacity to grow faster and form bigger white colonies. Absence of plasmid was confirmed by PCR for YopM / YopK. This pYV-derivative was renamed VpYV-.

### Animal housing, infection and *in vivo* analyses

Institut Pasteur animal facilities for animal housing are accredited by the French Ministry of Agriculture (accreditation B 75 15-01, 05/22/2008), in appliance of the French and European regulations on Laboratory Animals (EC Directive 86/609, French Law 2001-486 issued on June 6, 2001). The research protocol (N° 2013-0038) was approved by Institut Pasteur Ethics Committee for Animal Experimentation (CETEA) for the French Ministry of Research and was performed in compliance with the NIH Animal Welfare Insurance #A5476-01 issued on 02/07/2007.

Mouse infections were performed in a BSL3 animal facility [14]. Oral inoculation of bacteria by gavage with a cannula or on bread were performed as previously described; [25]. Immunodeficient mice MuMT [26] and RAGγc^-/-^ (a kind gift of Dr J. DiSanto [27] were bred in the Pasteur Institute animal facility. Corresponding C56BL/6J mice were obtained from Charles River France. Vaccination consisted of a single dose of bacteria in saline given either intragastrically (ig; 200 μl), subcutaneously in the ventral skin (sc; 50μl) or intradermally (id; 5 μl) in the ear tissue [28].

### Characterization of the immune response

Blood taken to obtain serum was collected from live animals by puncture of the maxillary artery with a Goldenrod lancet (Medipoint, USA) 3 weeks after vaccination or at the indicated time for long-term follow-up. IgG specific for *Yersinia* were quantified by ELISA as described before [29] [14], [30]. Microtiter plates (NUNC Maxisorp) were coated either with F1 antigen (10 μg/ml), or with 5 μg/ml of a sonicate of the *Y. pestis* CO92Δ*caf* strain grown at 37°C on LB agar, containing all *Y. pestis* antigens except F1. Antibody titers were calculated as the reciprocal of the lowest sample dilution giving a signal equal to two times the background. For the follow-up of immunity over time, heparinized blood was taken, and plasma was separated for IgG analysis, whereas leukocytes were isolated for *in-vitro* stimulation with 5 μg/ml of sterile soluble *Y. pestis* CO92 antigens obtained by bacteria sonication, as described previously [13].

### Preparation of *Yersinia* Whole-Cell Lysates and Supernatant

*Y. pseudotuberculosis* strains (VTnF1 and VpYV-) were grown in LB agar plates overnight at 28°C. *Yersinia pestis* CO92 pYV^+^ was growth in LB agar supplemented with Hemin 0.002%. From these plates, strains were grown in LB-MOX (LB Miller with 20mM of Sodium Oxalate and 20mM MgCl_2_). Bacterial suspensions (OD_600_ = 0.1) were incubated for 1 hour at 28°C and then moved to 37°C for 3 hours to reach exponential growth phase (VTnF1 and VpYV-) or overnight (*Y. pestis*). The fraction containing the T3SS effectors was obtained by centrifugation at 4500 rpm for 10 minutes and filtration of the conditioned culture supernatant on PES (0.22 μm) filters. Supernatant was gently collected and precipitated with TCA (Trichloroacetic acid, Sigma) ON at -20°C. Precipitated proteins were collected by centrifugation for 30 minutes at 4°C and washed with 400 μl of ice-cold acetone. After centrifuging again for 15 minutes, the pellet was air dried for 15 minutes.

*Y. pestis* whole cell lysates were prepared by lysing bacteria (grown at 37°C in LB-MOX) in Laemli buffer (Bio-Rad) containing ß-mercaptoethanol at 95°C for 10 minutes. The supernatants containing T3SS effectors were sterile filtered with PES filters (0.22 μm). Proteins were precipitated using Trichloroacetic acid (TCA) as described above. For loading on SDS-PAGE gels, T3SS effectors/proteins from supernatant were resuspended in Urea 8M and mixed with Laemmli buffer (Bio-Rad). OD_600_ of bacterial cultures was used to normalize the amount of sample for loading into the SDS-PAGE.

### SDS-PAGE and Western Blot Analysis

The different fractions of proteins were treated for SDS-PAGE and loaded in NuPAGE 4-12% (Invitrogen). Proteins were transferred to PVDF membranes using the Iblot® system and gel transfer stacks PVDF Mini (Invitrogen). Membranes were blocked with TBS-T (TBS-Tween 0,1 %) plus 5% milk (blocking buffer) and then incubated with different mouse sera diluted in TBS-T plus % milk. Western blots were revealed using Goat anti-mouse IgG (H+L) HRP conjugate (Bio-Rad) and membranes were washed between the different incubations for 2 cycles of 10 minutes with TBS-T. Finally, the substrate Pierce™ ECL Western Blotting Substrate (Thermo Fisher Scientific) was used to reveal chemiluminescent signal and read using an Amersham 680 imager.

### Evaluation of mouse protection against a challenge with *Y. pestis*

*Y. pestis* strains CO92 and its non-encapsulated (F1-negative) derivative CO92Δ*caf* [30] were grown at 28°C to prepare suspensions in saline solution for infections. To cause bubonic plague, subcutaneous injections (sc; 100 μl) were performed in the ventral skin with 10^3^ or 10^5^ CFU CO92 (10^2^xLD_50_ or 10^4^xLD_50_, respectively) or 10^5^ or 10^6^ CFU CO92Δ*caf* (10^3^xLD_50_ or 10^4^xLD_50_, respectively). To cause pneumonic plague, intranasal infection of anesthetized mice was performed as previously described [29] by instillation of 20 μl of bacterial suspension containing 10^5^ CFU (30xLD_50_) or 10^7^ CFU CO92 (3300xLD_50_). Survival was followed for 21 days.

### Host iron nutritional immunity

Protocols described by [31] were used. Briefly, mice either naïve or infected SC (back skin) 24h before with strains EV76 (*Y. pestis* ΔHPI) and VTnF1 and VpYV- (*Y. pseudotuberculosis* vaccine strains) were bled to produce serum. *Y. pestis* CO92 (10^5^ CFU/mL) was grown in these sera *in vitro* (300 μL cultures) and growth was monitored by plating aliquots. For protection assays, mice were co-injected with CO92 sc (100 CFU in ventral skin) and an attenuated strain (10^7^ CFU sc in back skin) and survival was monitored for 21 days.

### Statistical analyses

The Fisher exact test was used to compare mouse survival outcomes. The Mann-Whitney test was used to compare weight, antibody titers and IFNγ production. The Student’s T test was used to compare sera in *in-vitro* bacterial growth experiments.

## RESULTS

### Construction of VpYV-, an antibiotics-sensitive and pYV -cured derivative of strain VTnF1

To comply with “Good Manufacture Practices” (GMP), the four antibiotics-resistance cassettes: Kanamycin, Trimethoprim, Chloramphenicol and Spectinomycin / Streptomycin present in the genome of strain VTnF1 [30] were removed by means of allelic exchange, yielding strain VTnF1-S (S for susceptible).

The presence of any unwanted DNA (plasmidic, viral or other) accidentally inserted in the genome during engineering was excluded using Next Generation Sequencing of the strain’s genome. To exclude the possibility that elimination of the cassettes affected the strain’s immunogenicity, mice were inoculated orally with either VTnF1-R^-^ or VTnF1-S. Serum antibodies directed against F1 or other Yersinia antigens were equivalent for the two strains (Supplementary Figure S1A) and VTnF1-S induced protection against plague as efficiently (Supplementary Figure S1B).

As previously reported for VTnF1-R, vaccinated mice presented a transient weight loss indicating a systemic reaction (Supplementary. Figure S2). We hypothesized that the presence of the T3SS (minus YopK) in the strain could contribute to this residual virulence. A completely T3SS deficient derivative of VTnF1-S was obtained by curing the pYV plasmid and was renamed VpYV-. By SDS-Page electrophoresis, we could verify that production of effector Yops by bacteria in low Calcium conditions (LCR) was higher in VTnF1 compared to it parental strain (Figure 3A) due to the YopK mutation [12] and abolished in VpYV-.

### *In-vivo* virulence and dissemination of the VpYV-strain

Live attenuated vaccines can exert severe secondary effects on immunocompromised subjects, and such a vaccination is not recommended for them. When inoculated orally to normal laboratory mice (OF1) at doses of 10^8^ CFU (a total of 124 mice were tested), 10^9^ CFU (N=44) or 10^10^ CFU (the maximum which was possible to test; N=28), VpYV-induced no clinical signs such as ruffled fur or weight loss (Supplementary Figure S2). Two immunocompromised mice lines : MuMT mice producing no antibody (B cell deficient) and RAGγc^-/-^ mice (multiple T/B/NK cells deficiency) were compared to genetic background-matched immunocompetent mice (C57BL/6) for their ability to control the live VpYV-strain given orally (Figure 1). Whereas a single oral inoculation of VTnF1 was fatal to most MuMT and all RAGγc^-/-^ mice, VpYV-did not cause lethality in these immunocompromised mice, demonstrating its strongly enhanced attenuation.

**Figure 1:**
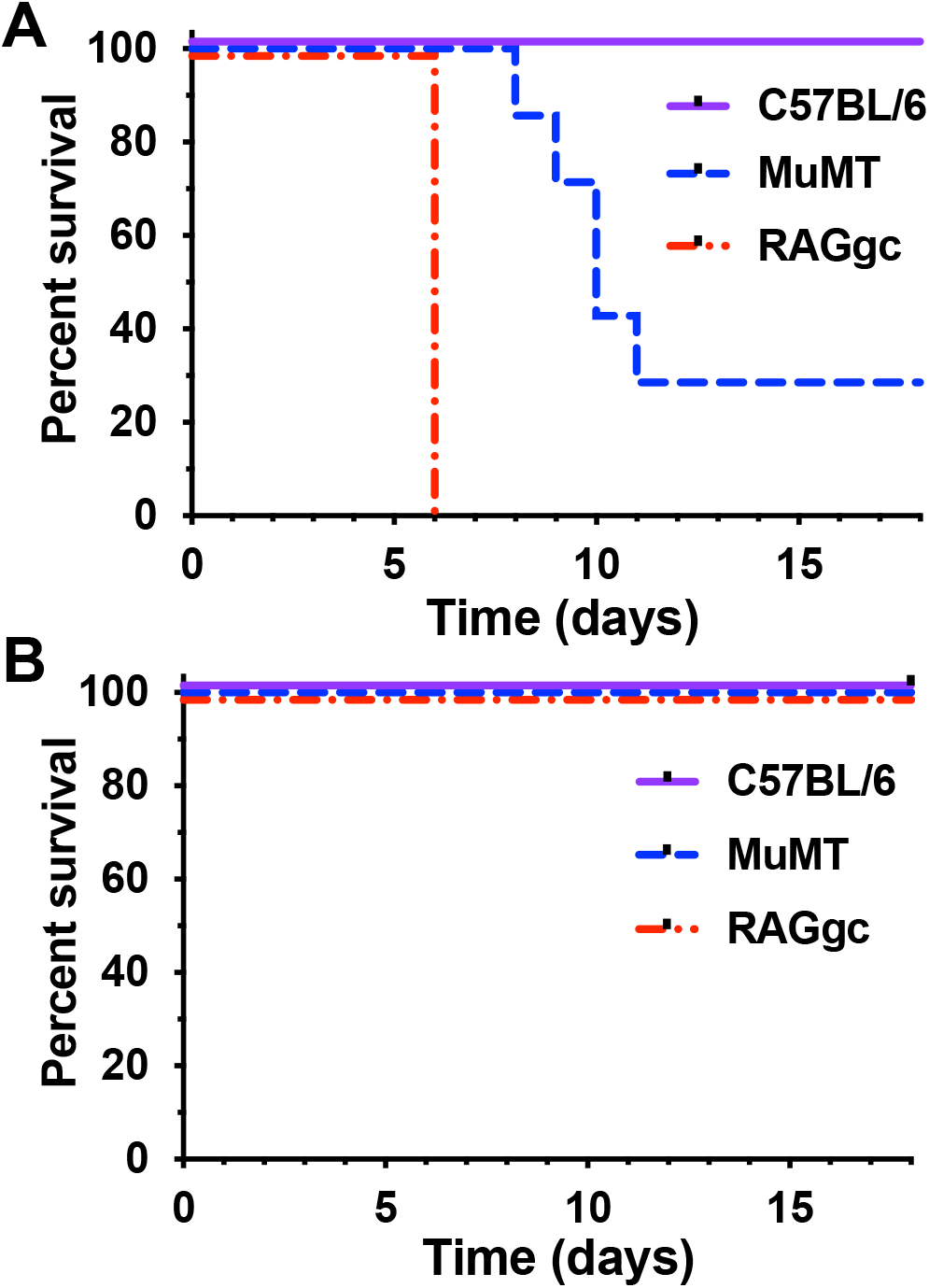
Survival of immune-depressed mice to oral vaccination against plague with VTnF1 or its virulence plasmid-cured derivative. Mice (7 per group) received one dose (10^8^ CFU) of VTnF1 (A) or the plasmid-cured derivative VpYV- (B) intragastrically and survival was followed for 18 days.

The capacity of VpYV-to disseminate *in vivo* was evaluated by measuring the bacterial load in tissues of mice at different time points after vaccination. VpYV-colonized the Peyer’s patches as efficiently as the plasmid-positive VTnF1 [32] and then accumulated in mesenteric lymph nodes (Figure 2). Its progression mostly halts there and progression to the liver and spleen was weak, indicating a capacity to follow the lymphoid network, but a reduced ability to gain access to blood and blood-filtering organs (spleen & liver). Bacteria were also searched for in the cecum 34 days after vaccination because virulent *Y. pseudotuberculosis* can persist there for long times and cause abscesses [33]. No bacteria were found, in agreement with the view that this ability requires more virulence.

**Figure 2:**
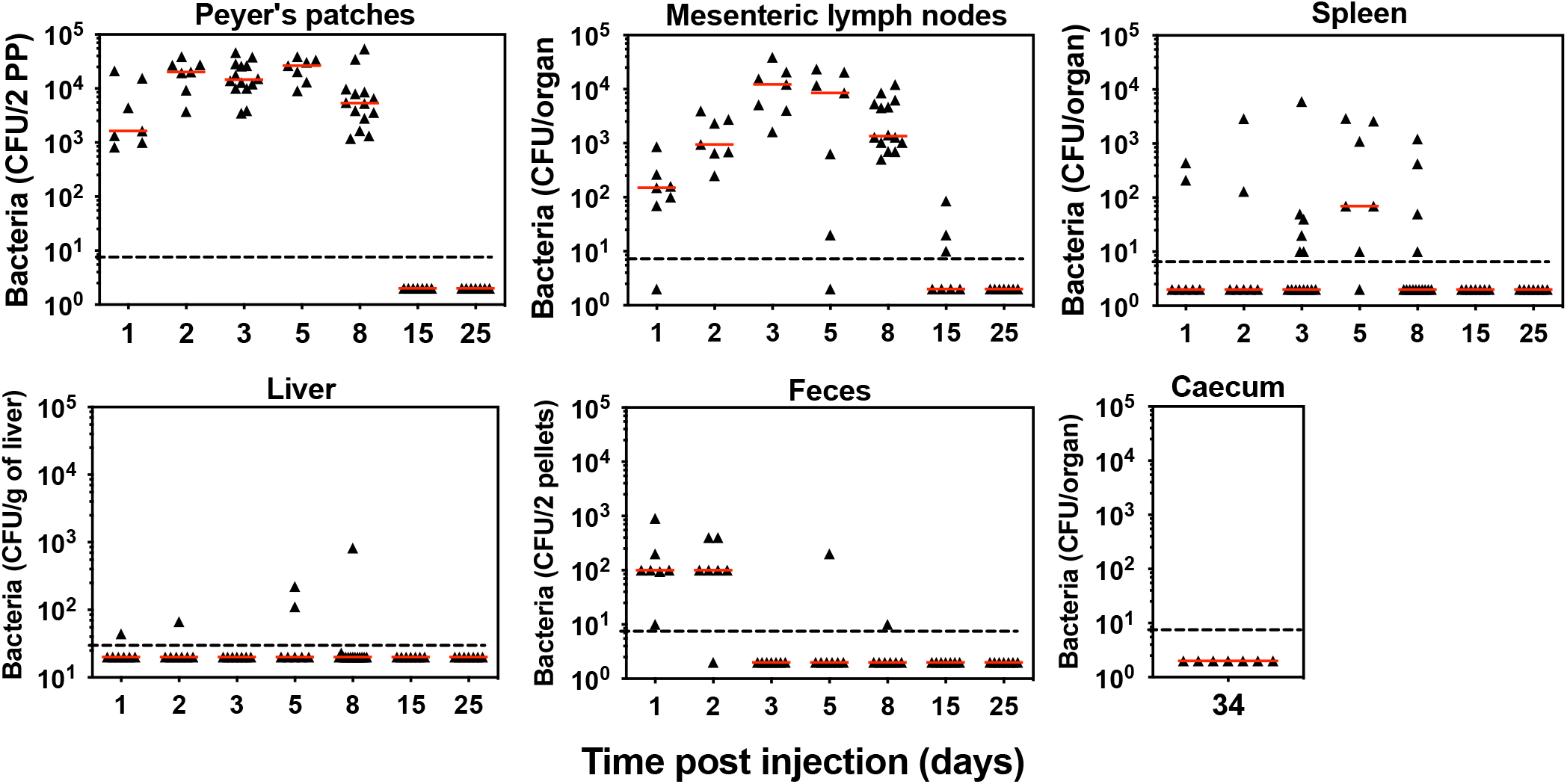
*In vivo* dissemination of the VpYV-strain in orally vaccinated mice. A single dose of strain VpYV-(10^8^ CFU) was inoculated orally to groups of 7-14 mice. Mice were humanely sacrificed for organ collection at predetermined times to evaluate the bacterial loads in feces and indicated organs. Samples were minced and dilutions were plated on selective agar plates to count colonies, with a detection limit of 10 CFU/sample. Shown are individual results and the median (horizontal bar).

### Immunogenicity and protective value of the VpYV-strain against bubonic and pneumonic plague

The immunogenicity of VpYV-was evaluated by measuring serum IgG levels from mice receiving various doses orally. Titers IgG against the F1 antigen were comparable to those induced by VTnF1, whereas VpYV-failed to trigger as high IgG titers against non-F1 *Yersinia* antigens (Figure. 3B), as confirmed by a lower number of *Y. pestis* antigenic targets recognized by immune sera by Western blot (Figure 3C) When re-examined 4 month after vaccination with VpYV-, IgG titers were still high, although IgG against F1 had started to decrease (Figure 3D).

An IFNγ release assay using leukocytes to test the cellular side of the immune response revealed that most mice developed a cellular response against the vaccine VpYV-(Figure 3E). IFNγ levels did not weaken after 4 months, however levels were lower than those observed previously with the VTnF1 strain [13], indicating a lower immunogenicity of the VpYV-strain.

**Figure 3:**
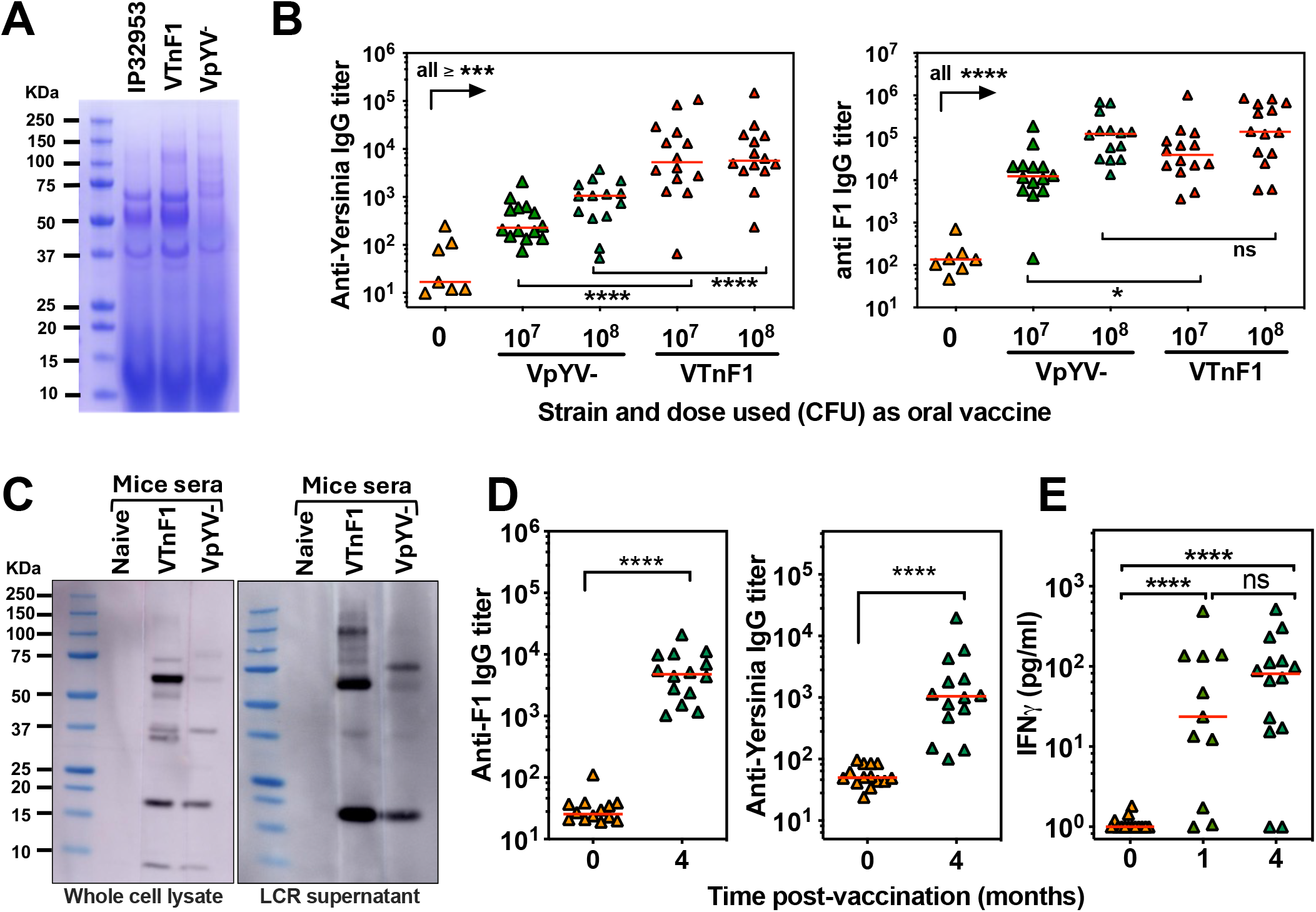
Immune response induced by VpYV- oral vaccination. (A) Secretion of T3SS effectors in the different vaccine strains (VTnF1 and VpYV-) and their background strain *Yersinia pseudotuberculosis* IP32953. (B) The plasmid-cured VpYV-strain was given orally to mice at the indicated doses and was compared to the parent VTnF1 strain. Serum IgG production was quantified 3 weeks post vaccination using an ELISA measuring IgG directed against *Y. pestis* antigens other than F1 or purified F1 antigen. (C) Sera of mice vaccinated with VTnF1 or VpYV-(10^8^ CFU orally) or not vaccinated were used to reveal target *Y. pestis* antigens (37°C culture and low calcium) present in whole cell lysates and supernatant blotted after SDS-Page separation. (D) IgG were quantified as in (B) in sera taken before or 4 months after oral vaccination with VpYV-. (E) Leukocytes from mice vaccinated 1 month before were stimulated *in vitro* with 5 μg/ml of sterile soluble *Y. pestis* CO92 antigens. IFNγ was measured by ELISA in day 3 culture supernatants. Shown are results from 14 vaccinated animals and 7 (B) or 14 (D-E) naïve ones. Horizontal red lines indicate group medians. The Mann-Whitney test was used for statistical analysis: *: p<0.05; **: p<0.01; ***: p<0.001; ****: p<0.0001. ns: not significant.

Protection against plague was tested against the bubonic and pneumonic forms of the disease. Against bubonic plague, oral inoculation of VpYV-protected 100% of the mice, even when a 100 x higher challenge dose (Table 1). This protective immunity needed 9 days to fully develop (Supplementary Figure S3), and still saved 64% of the mice 6 months later (Table 1). Against pneumonic plague, VpYV-could protect 79% of mice and this fell to 28% after 6 months.

**Table 1:**
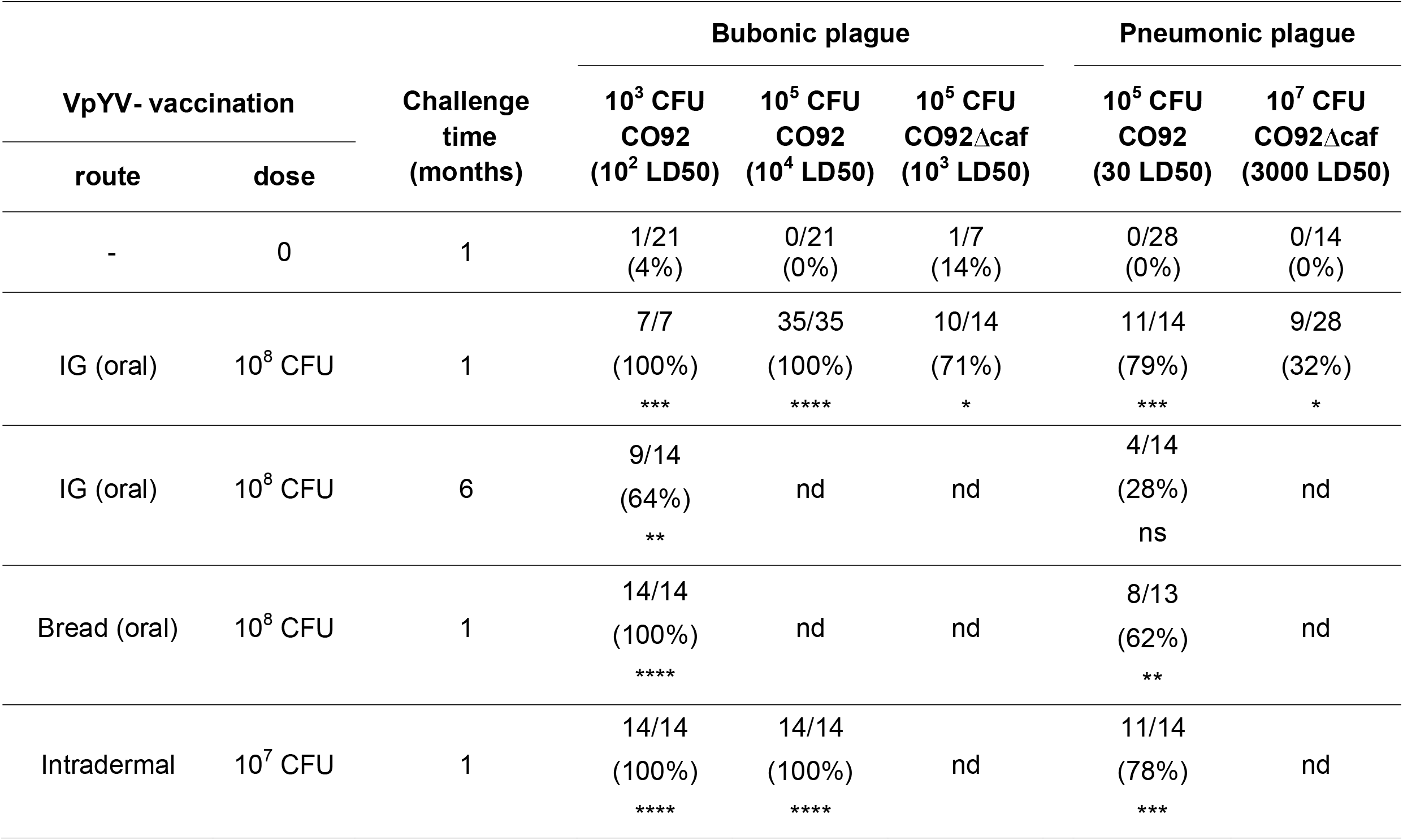
Protection of mice vaccinated with VpYV-against bubonic and pneumonic plague Mice having received a single dose of VTnF1 were challenged 4 weeks later by infection with *Y. pestis* CO92 s.c. (bubonic plague) or i.n. (pneumonic plague). Survival was recorded daily for 21 days. Protection significance was tested using Fisher’s two-tailed exact test, compared to naïve mice of the same experiment : *: p<0.05; **: p<0.01; ***: p<0.001; ****: p<0.0001

These results were obtained by intragastric inoculation of bacteria to mice with a cannula, before we developed an oral infection model based on volunteer eating of bread loaded with bacteria as a more innocuous method [25]. Both methods in fact gave similar results: VpYV-triggered a comparable antibody response and conferred equivalent protection against bubonic and pneumonic plague (Supplementary Figure S4A&B and Table 1). Finally, intradermal vaccination was also tested as an alternative to oral vaccination [34] and was as effective to induce specific IgG (Supplementary Figure S5) and was able to protect against bubonic plague as when the oral route was used (Table 1).

Also, VTnF1 had the remarkable ability to protect against plague caused by a F1-negative *Y. pestis* variant (CO92Δ*caf*; [30]). When tested in the same conditions, VpYV-was 30 % less efficient against bubonic plague and much less against pneumonic plague (Table 1), revealing that the pYV plasmid and the tools it encodes are mandatory to obtain full protection against F1-negative *Y. pestis*.

### ‘Host iron nutritional immunity against plague’ can be induced by the attenuated VTnF1 *Y. pseudotuberculosis* strain but requires its pYV plasmid

It was recently reported that distal co-injection of the live attenuated *Y. pestis* EV76 strain conferred protection against infection by a wild-type strain in mice, suggesting a possible urgence therapeutic means [31]. The antibacterial activity was mediated by a fast production of host heme- and iron-binding proteins, so that *Y. pestis* growth *in vitro* was reduced in sera from mice recently inoculated with EV76. We tested whether this ‘iron nutritional activity’ could be induced by our vaccine strains, either VTnF1 or VpYV-, which (like EV76) do not have the HPI [30]. Sera from mice infected with EV76 or strains VTnF1 and VpYV-was able to hamper growth *in vitro* as compared to naïve serum (Figure 4A), although VpYV-’s effect was intermediate (Figure 4A). This confirmed the observations of Zauberman *et al*. with EV76 and extended it to attenuated *Y. pseudotuberculosis* HPI^-^ strains.

**Figure 4:**
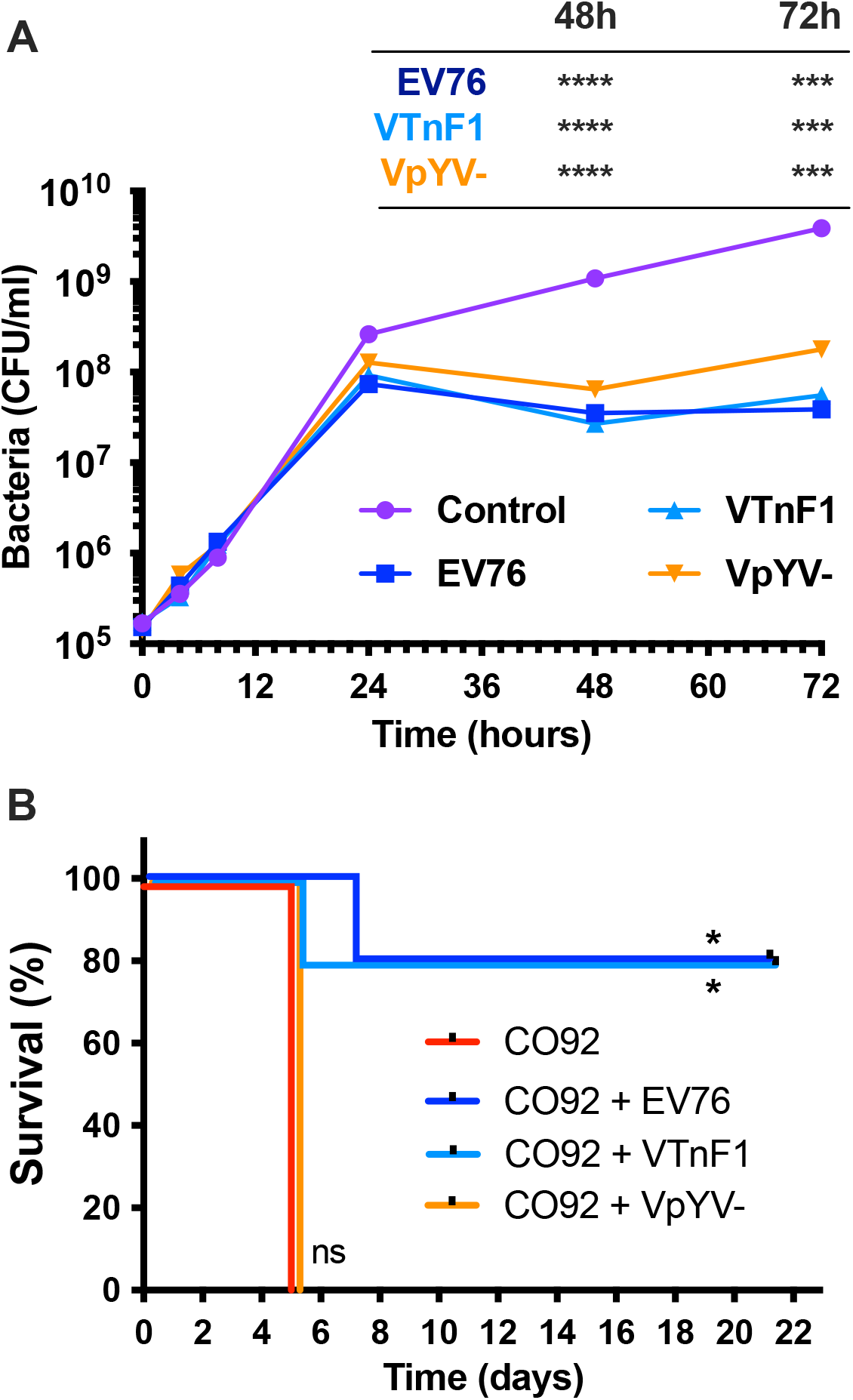
Host iron nutritional immunity induced by *Y. pseudotuberculosis* vaccine strains. (A) Sera was obtained from mice immunized with either EV76, VTnF1 or VpYV-strains or naïve mice, and was used to grow *Y. pestis in vitro* and follow bacteria growth. Shown are values from one representative experiment out of 2. The t test was used to compare non-immune serum to immune ones : ***: p<0.001; ****: p<0.0001. (B) Mice were infected subcutaneously with *Y. pestis* and co-infected using the same route but at a distal site with 10^7^ CFU of EV76, VTnF1 or VpYV- and survival was followed. The Fisher Exact tests was used: *: p<0.05.

The protective value of co-infection *in-vivo* was tested by co-infecting mice with *Y. pestis* CO92 with each of the HPI^-^ strains, EV76, VTnF1 and VpYV-. Co-injection of EV76 protected most mice against plague (Figure 4B), and so did VTnF1. However, strain VpYV-failed to protect mice. Altogether, this indicated that an attenuated *Y. pseudotuberculosis* strain was able to exert the protective ‘iron nutritional activity’ against plague as efficiently as *Y. pestis* EV76 but required the pYV plasmid to exert this activity.

## DISCUSSION

The goal of vaccination is to reduce the risk of getting a disease by educating the body’s immune system to recognize the invasive germ, such as viruses or bacteria. Live attenuated vaccines therefore need a fine tuning between their ability to wake the system (immunogenicity) and the risk of causing damage (virulence). We here evaluated the possibility of removing completely the T3SS in a live attenuated plague vaccine to obtain the best possible attenuation, and report on the consequences on its efficacy. Overall, the price to pay for more attenuation is a lesser immunogenicity, with less but still effective protection against plague, showing that the T3SS has a clear contribution to vaccine’s characteristics.

VpYV-has an ability to disseminate into the intestine’s lymphoid tissues (Peyers’ patches and mesenteric Lymph nodes), but comparison with VTnF1 shows that it yields lower bacterial loads in these lymphoid structures [32]. In addition, it has a clear defect in penetrating deeper tissues (spleen, liver), blood filtrating organs thought to be contaminated via blood during yersiniosis. Therefore, our observations indicate that the pYV-cured strain is much less able to reach the blood, and that means less tissue destruction in lymph nodes. In agreement, a Yop-deficient *Y. pseudotuberculosis* was defective in intestinal tissue colonization [35]. Balada-Llasat previously demonstrated that *Y. pseudotuberculosis* mutants lacking the pYV survived as well as or better than wild-type (WT) ones in the mesenteric lymph nodes (MLN), homing inside B-T rich regions where they are protected [36]. This tropism could be a reason for the fairly good immunogenicity of VpYV-regardless of its inability to settle in the spleen. Similarly, a pYV-cured *Y. pseudotuberculosis* YPIII strain given orally provided protective immunity against pseudotuberculosis when given orally [37]. Interestingly, that strain weakly developed in the spleen when injected iv, but partially protected against bubonic plague [21], suggesting it could have found there a T-B rich zone niche as well.

Studies of *Y. pestis* attenuated by curing its virulence plasmid yielded similar observations. The historical avirulent *Y. pestis* vaccine strain named Tjiwidej, developed by Haffkine in 1901 in Bombay was cured of its pYV/pCD1 plasmid, and was less protective than the pYV-positive strain EV76 produced soon after by Girard and Robic [17]. That a pYV-cured *Y. pestis* vaccine did not protect well may be due to the fact that *Y. pestis* is a stealth pathogen, which is much less detected by innate immunity receptors, due in part to differences in its LPS structure and absence of flagellin [17]. That EV76 induce better protection may result from the presence of pYV, causing tissue damage compensating for the weak detection of the molecular patterns.

Less stealth than *Y. pestis*, the *Y. pseudotuberculosis* on which VTnF1 and VpYV are built have a normal LPS and a flagellum, and thus are easily detected by host TLR2/4 and TLR5. The T3SS partly counteracts this innate response by targeting the host inflammasomes, intracellular polymeric platforms of the innate immune response which assemble in response to signals from TLRs, NLRs, and other innate sensors detecting LPS, flagellin and other Pathogen-Associated Patterns (PAMPs; [38], [39]). They lead to activation of the caspase-1 signaling cascade giving rise to the release of pro-inflammatory factors which include cytokines (IL-1ß, IL-18) and alarmins (IL-1α, HMGB1), as well as pyroptotic cell death [40]. By causing inflammation, they play a central role in the elicitation of an adaptive response as well, and their activation is therefore important to a vaccination process [41]. The injected T3SS effectors generally targets multiple inflammasomes and combine both activator and inhibitory effects. While some pathogenic bacteria take advantage of their activation (e.g. *L. monocytogenes, H. pylori*), T3SS Yop effectors of pathogenic *Yersinia* sp. exert effects in both directions, but overall subvert inflammasomes to prevent inflammation [40]. Therefore, removing the whole T3SS has both positive and negative effects on inflammasome activation. When the VTnF1 strain was created, we thought that Yops would be good targets of immunity, and that eliminating the T3SS apparatus could have caused a too fast elimination of the vaccine *in vivo*, and hence would drive a weak immune response. Deletion of the *yopK* gene attenuated VTnF1 because YopK controls the flux of effector Yops [12] and is itself an inflammasome inhibitor [42], but it did not fully make the strain avirulent.

By eliminating the whole T3SS in VpYV-we took the bet that it could prevent the remaining inhibitory effects of Yops, avoid inhibition of some inflammasomes, and consequently favor immunity against the other antigenic targets, at the risk that the bacilli could be faster eliminated. Indeed, attenuation is stronger, the strain disseminates less and persists less, but also induces a less intense immune response and les protection. The T3S was therefore a key component of VTnF1’s efficacy.

On the other hand, immunity against the T3S can be induced and is desirable. Many molecules of the TTSS machinery, including Yops, LcrV and Ysc elements of the injectisome, are immunogenic. Although several Yops do not induce a protective immune response when used alone as vaccine [43], protection was obtained by vaccinating with structural elements of the injectisome, such as the tip LcrV, the YopBDE translocon complex [44] and the abundant needle component YscF [45]. Notably, the injectisome tip LcrV is an excellent target of immunity which was included in some recent molecular candidate plague vaccines [46]. However, VTnF1 had failed to induce a response against the LcrV antigen [32], similarly to a *Y. pestis*-based live vaccine [47], so its absence from VpYV cannot have affected the strain’s immunogenicity. Finally, a likely disadvantage of losing the T3S results from the fact that it favors a host Th17 cellular response [48], with it hallmark IL-17 production being important against pneumonic plague [49]. This may explain why protection against this form of plague was not as good with VpYV-as with VTnF1.

Protection against F1-negative *Y. pestis* is a goal imposed by the idea that such a variant would be a better bioterrorist weapon, unseen by diagnostic dipsticks detecting F1, and in part resistant to vaccines built on F1 + LcrV antigens [46]. Indeed, at least three major laboratories in the world are developing molecular plague vaccines based on these antigens, considered as the best targets at *Y. pestis*’ surface [46]. As previously reported and described here, VTnF1 and VpYV-induce a clear response against F1 and multiple other antigens. However, VTnF1 does not induce an antibody response against LcrV, although it confers protection against an F1-negative *Y. pestis* (CO92Δ*caf*; [52]), indicating that the other antigens, including Yops, are efficient targets. Instead, the absence of response against LcrV, the absence of Yops and the uselessness of F1 against CO92Δ*caf* may explain the difficulty of VpYV-to protect against an F1-*Y. pestis*.

## Supporting information

2 supplemental tables plus 5 supplemental figures

## Acknowledgements

This work was supported by an ANR “Emergence 2012” grant (N°EMMA-2012-0011-01). The Yersinia Research Unit is a member of the LabEX IBEID (ANR LBX-62 IBEID).

