## Supplementary material for "Contribution of the Type 3 Secretion System to immunogenicity of a live *Yersinia pseudotuberculosis* plague vaccine": 2 supplemental tables plus 5 supplemental figures

**SUPPLEMENTARY DATA**

**Supplementary Table S1: Strains and plasmids used in this study**

| **Strains/plasmids** | **Genetic characteristics** | **Antibiotic markers** | **ref** |
| --- | --- | --- | --- |
| **Strains** | | | |
| *E. coli* BW19610 | λ*pir* | - | 1 |
| *E. coli* SM10 | λ*pir thi thr leu tonA lacY supE recA::*RP4-2-Tc::Mu(Km) | Km | 2 |
| Y*. pseudotuberculosis* IP32953 modified =  VTnF1 | ΔHPI, *psaA*, *yopK* ::Tn*7*-*caf*  pYV+ | Km, Tmp, Spec, Cm | 3 |
| *Y. pseudotuberculosis*  VTnF1-CmS | ΔHPI, *psaA*, *yopK* ::Tn*7*-*caf*  pYV+ | Km, Tmp, Spec | This study |
| *Y. pseudotuberculosis*  VTnF1-KmS | id | Tmp, Spec | This study |
| *Y. pseudotuberculosis*  VTnF1-TmpS | id | Spec | This study |
| *Y. pseudotuberculosis*  VTnF1-S | id | - | This study |
| *Y. pseudotuberculosis*  VpYV- | ΔHPI, *psaA*, *yopK* ::Tn7-*caf*  pYV- | - | This study |
| **Plasmids** | | | |
| pFLP3 | FRT resolvase, *bla*, *sacB* | Amp | 4 |
| pCVD442 | R6K *pir* dependent replication, *mobRP4*, *bla*, *sacB* | Amp | 5 |
| pCVD442-up/down HPI | HPI-Upstream and downstream regions cloned in the SalI / SphI and SphI / SmaI sites, respectively | Amp | This study |
| pCVD442-up/down *psaA* | *psaA*-Upstream and downstream regions cloned in the SalI / SphI and SphI / SmaI sites, respectively | Amp | This study |
| pCVD442-up/down *yopK* | *yopK*-Upstream and downstream regions cloned in the SalI / SphI and SphI / SmaI sites, respectively | Amp | This study |

**Supplementary Table S2: Primers used in this study**

| **Name** | **Target** | **Sequence**  **(5’-3’)** |
| --- | --- | --- |
| *psaA*-up-forward | upstream region of *psaA* | ACGCgtcgacGGGTACAAGGAGAACATATCCATACG  SalI |
| *psaA*-up-reverse |  | ACATgcatgcGAGAACAGTCTCCATTAAATGTAATAATTGC  SphI |
| *psaA*-down-forward | downstream region of *psaA* | ACATgcatgcACGAGTAATCATAGAACATGATGGC  SphI |
| *psaA*-down-reverse |  | TCCcccgggCGATCTTCAGGGAATTTTCCACCG  SmaI |
| HPI-up-forward | upstream region of HPI | ACGCgtcgacCATTGCCCGTTTGGTTGTTGATAACAGCG  SalI |
| HPI-up-reverse |  | ACATgcatgcAAGAACCCTGCCTAGGTAACTAAGC  SphI |
| HPI-down-forward | downstream region of HPI | ACATgcatgcTGGCTCCTCTGACTGGACTCG  SphI |
| HPI-down-reverse |  | TCCcccgggCCGAACATTGTCGCTCAGCC  SmaI |
| *yopK*-up-forward | upstream region of *yopK* | ACGCgtcgacGGTGTGCGCCAATCACCCCTTCG  SalI |
| *yopK* -up-reverse |  | ACATgcatgcAGTTACTACTCCCAAATTTACTTTA  SphI |
| *yopK* -down-forward | downstream region of *yopK* | ACATgcatgcAGCTATATTAAAGAGTTTGGGAT  SphI |
| *yopK* -down-reverse |  | TCCcccgggCCACAGCCGGAGAGGACTAATAC  SmaI |

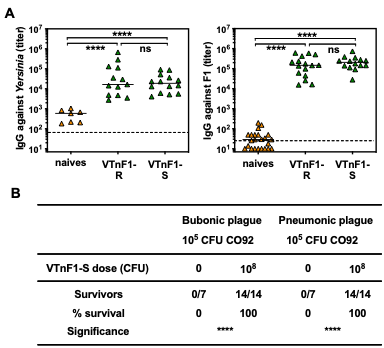

**Supplementary Figure S1: Humoral immune response and protection against plague induced by VTnF1-R or the antibiotics cassettes-free VTnF1-S strain**

Mice were vaccinated ig with the original VTnF1(-R) strain or the antibiotics-resistance cassettes free VTnF1-S version (10^8^ CFU ig; 14 mice) or were not vaccinated (naïves; 7 mice). (A): Sera were collected 3 weeks after vaccination and IgG specific for *Yersinia* antigens or purified F1 antigen were determined by ELISA. Shown are the limit of detection (dotted lines) and the median (short lines). (B): 2 groups of mice vaccinated with VTnF1-S were challenged 4 weeks later by infection with *Y. pestis* CO92 s.c. (bubonic plague) or i.n. (pneumonic plague). Survival was recorded daily for 21 days. Protection significance was tested using Fisher’s two-tailed exact test, ***: p<0.001; ****: p<0.0001

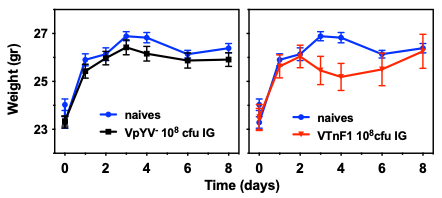

**Supplementary Figure S2: Weight monitoring after oral vaccination with the VTnF1-S or VpYV- strains.**

Mice were vaccinated ig with the VTnF1-S strain or the plasmid-cured VpYV- strain (10^8^ CFU ig ; 14 mice) or were not vaccinated (14 mice). Individual weight was followed daily during the week after vaccination Shown is the mean ± sem.

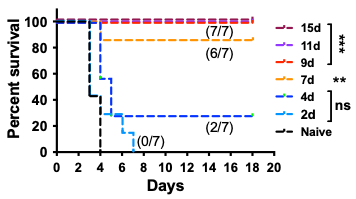

**Supplementary Figure S3: Kinetics of protective immunity against bubonic plague in mice vaccinated orally with VpYV-.**

Groups of mice (n=7) received one oral dose of VpYV- strain (10^8^ CFU) and were infected at increasing times after with *Y. pestis* CO92 injected sc (10^5^ CFU; bubonic plague). Survival was followed for 18 days. The Fisher Exact test was used for statistical analysis of group survival compared to the naïves group : **: p<0.01; ***: p<0.001. ns: not significant.

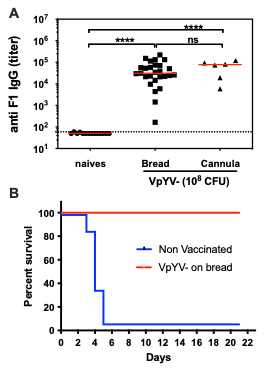

**Supplementary Figure S4: Vaccination with VpYV- using the bread-feeding method triggers an equivalent immune response and protection against plague.**

The VpYV- vaccine (10^7^ CFU) was given to mice either with a cannula or on bread for volunteer eating by animals. Two mice did not eat the complete bread piece and developed a lower antibody response. (A) Sera were collected 3 weeks after vaccination and IgG specific for Yersinia antigens or purified F1 antigen were determined by Elisa. Shown are the limit of detection (dotted lines) and the median (short red lines). (B) Mice were challenged 4 weeks later by infection with *Y. pestis* CO92 s.c. (bubonic plague). The Mann-Whitney test was used for statistical analysis: ****: p<0.0001. ns: not significant.

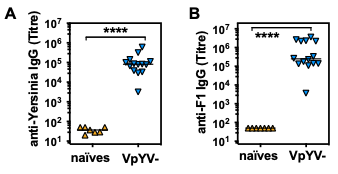

**Supplementary Figure S5: Immune response of mice vaccinated with VpYV- by the intradermal route.**

Groups of mice were injected ID with strain VpYV- or were not vaccinated (naïves). Blood was taken 3 weeks pi and serum IgG against purified F1, or other antigens were measured by ELISA.
